# Reticulate evolution in a neutral model: speciation, extinctions, and hybridizations

**DOI:** 10.1101/2021.07.19.452966

**Authors:** Larissa Lubiana Botelho, Flavia Maria Darcie Marquitti, Marcus Aloizio Martinez de Aguiar

## Abstract

Evolution is usually pictured as a tree where ancient species branch into new ones and eventually disappear. In this simplified view, the balance between speciation and extinction fully determines the diversity of life. Hybridization, however, introduces another level of complexity, allowing neighboring branches of the tree to interact, mixing their genetic content. This generates further diversity leading to reticulated phylogenetic trees. In this paper we study processes of speciation, extinction and hybridization using a genetically and spatially explicit neutral model of diversification. Speciation, extinction and hybridization events are tracked throughout the evolutionary process leading to complete and exact phylogenetic trees. We found that genome size played a key role in these processes, increasing the extinction rate and decreasing the hybridization rate. In our simulations, hybridization after one speciation event occurred throughout the evolutionary process but hybridization after two speciation events was only observed during the initial radiation. Most hybridization occurred between relatively abundant species, discarding lack of sexual partners or small population sizes as potential causes. We found that hybridization occurred mostly because of opportunity (genetic similarity and spatial proximity) between recently branched species, when the number of accumulated mutations is not yet too large.

## Introduction

The evolutionary process cannot always be represented by a tree like structure where species, once formed, can only branch again into new species or go extinct. Hybridization, through the breakdown of reproductive barriers, lineage fusion and speciation reversal, can merge branches, introducing loops, or reticulation, in the phylogenetic structure (Kleindorfer et al., 2014; Garrick et al., 2014; MacLeod et al., 2015; Rudman and Schluter, 2016; Kearns et al., 2018; Pyron et al., 2020).

Phylogenies that do not take into consideration processes that lead to reticulation, like hybridization, can lead to the construction of incompatible trees that are not verisimilar representations of the real tree (Zhang et al., 2021). Detecting hybridization, however, can be challenging. Purebreds, for example, can quickly be replaced by a group of admixed individuals with few signatures of its evolutionary history (Garrick et al., 2014). Also, newly separated species with weak reproductive barriers can interbreed again for short periods of time before eventually splitting into well separated species, leaving no records of the hybrids (Seehausen, 2004). However molecular tools, such as mitochondrial DNA analysis, are allowing the detection of introgressive hybridization in extant species. Galapagos giant tortoises from the island of Isabela (Garrick et al., 2014) and marine iguanas from the San Cristóbal island (MacLeod et al., 2015) are very well studied species that have undergone fusion processes. Moreover, even after reproductive isolation has been established, speciation reversal might still occur, as has been found in the North American Common Ravens (Kearns et al., 2018). Speciation has also been undone among finches of the Floreana Island and it seems to be responsible for the disappearance of the large tree finch *Ca-marhynchus psittacula* and it is likely that other two species of finches (*C. pauper* and *C. parvulus*) are merging into one (Kleindorfer et al., 2014). Recently, the merging of two *Geospiza* species (*G. fortis* and *G. scandens*) has been happening and it is observed since 1982. The forecast, according to the rates of introgres-sion, is that they complete the speciation in reverse between 2049 (genetically) and 2057 (morphologically) (Grant et al., 2004; Grant and Grant, 2014).

Usually, hybridization is referred to two opposite effects: first as an important process by which new pathways to genetic diversity and novelties arise and, second, as a mechanism by which species diversity is diminished. Hybridization increases the number of species if there is no backcrossing to the parents and if speciation happens after as a consequence, for instance, of advantageous novelties or by fast reproductive isolation (Seehausen, 2004; Abbott et al., 2013; Lamichhaney et al., 2018). On the other hand, hybridization can cause loss of diversity by the extinction of species (through genetic swamping and demographic swamping), lineage fusion, speciation reversal and by the breakdown of reproductive barriers (Todesco et al., 2016). In this study we focused on hybridization as a mechanism through which diversity is diminished. Many factors can boost the occurrence of hybridization events: the scarcity of sexual mates (Willis et al., 2011), the recognition failure of co-specific partners (e.g., misimprinting) (Grant and Grant, 1997, 2014), and also fast environmental changes and anthropogenic changes, such as climate change and habitat loss (Seehausen et al., 2008; Kleindorfer et al., 2014), are pointed as important forces for increasing reproduction between individuals of different species. However, the importance of hybridization in decreasing diversity may be underrated because of the difficult to detect it in biological data and also because many of them can happen only for a few generations.

In this study we present an extension of a spatially explicit evolutionary model in a neutral scenario (de Aguiar et al., 2009; de Aguiar, 2017; Costa et al., 2019) and use it to characterize the processes of hybridization and extinction. In this model, in which species emerge in response to genetic and spatial restrictions imposed on reproduction, processes that decrease diversity (extinction and hybridization) are also emergent events. We start with a genetically uniform population, which radiates into many species until it reaches an equilibration of the species diversity (see Fig. S1 in the SM showing the number of extant species across generations). To characterize the different processes that occur during its evolution we keep track, at every generation, of all events of speciation, extinction and hybridization. This allows us to compute true and complete phylogenies, from its root at *T* = 0 to the leafs at present time, containing all events that ever occured in that population. This offers a theoretical methodology that avoids the common loss of evolutionary information caused by gene flow (Zhang et al., 2021). Previous theoretical studies have focused on hybridization (and on extinction in some cases) using non-neutral models (McCarthy et al., 1995; Buerkle et al., 2000; Duenez-Guzman et al., 2009; Kagawa and Takimoto, 2018). In this work we use a neutral model in which individuals have no advantages or disadvantages in their reproduction, or in their death probabilities, and mating is restricted by spatial proximity and genetic similarity (only individuals belonging to the same species can mate) but is otherwise random.

We focus our analysis on three points concerning the events of true extinctions and hybridization: (i) when they occur in the evolutionary history; (ii) what are species abundances at the moment of the event and; (iii) what is the life time of the species up to the moment of the event. We also show that these features are strongly affected by the amount of information that are important for the genetic isolation. We believe that the characterization of these events in a neutral scenario can shed light on how hybridization events contribute to the evolutionary history.

## Methods

### Evolutionary model

Our simulations were performed with the individual based model (IBM) of speciation proposed in Costa et al. (2019), which is an adaptation of the model of de Aguiar et al. (2009). The parameters used in this work are similar to those used in de Aguiar et al. (2009), which generated patterns of diversity compatible with empirical data. One of these parameters is the genome size *B*. We remark, however, that *B* should not be directly compared with number of genes or base pairs in real genomes. The number of loci should rather be interpreted as the amount of inheritable information carried by the individual, that we roughly classify as small (*B* = 150), medium (*B* = 1500) and large (*B* = 15000). In the simulations, population sizes are small, otherwise computational times becomes too large. The relation between the model parameters and the viability of speciation are discussed in (Baptestini et al., 2013).

We simulate the evolution of a population of *M* haploid individuals, randomly distributed in a spatial square area consisting of *L × L* discrete lattice sites with periodic boundary conditions. Individuals are assigned a genome represented by a binary string of size 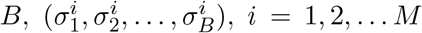, where each *locus k* of an individual *i*, 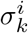, can assume the allele values 0 or 1. Their spatial position is specified by the coordinates (*x*_*i*_, *y*_*i*_) on the lattice plane. The genetic distance *d*^(*i,j*)^ between two individuals *i* and *j* is the Hamming distance between the corresponding sequences and measures the number of genes bearing different alleles:

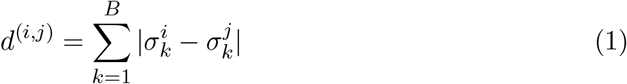

The spatial distance *s*^(*i,j*)^ between individuals *i* and *j* can be computed as:

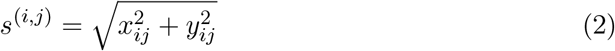

where *x*_*ij*_ and *y*_*ij*_ are distances between the individuals *i* and *j* in the *x* and *y* directions, taking into account the periodic boundary conditions.

Reproduction is sexual and time is counted in number of generations, which are non-overlapping. Mate choice is restricted by spatial and genetic distances. The spatial radius *S* defines the *mating range*, a circular neighborhood centered on the *focal* individual, where it looks for potential mates, and *G* specifies the *genetic threshold of reproductive isolation*, which is the maximum number of genetic differences that defines compatibility. Therefore, the focal individual *i* mates only with a compatible individual *j* (*s*^(*i,j*)^ ≤ *S* and *d*(*i, j*) ≤ *G*), that is chosen randomly with uniform probability. In the simulations, we used *G* = 0.05*B* and different sizes of genome: *B* = *{*150; 1500; 15000*}*. Each individual has a chance of reproducing, but there is a probability *Q* that it will not do so. In that case, another individual from its mating range is randomly chosen to reproduce in its place, keeping the population size constant. In our simulations, we set *Q* = 0.37 ≈ *e*^−1^, which corresponds approximately to the probability that an individual is not selected in *M* trials with replacement, in accordance with the Derrida-Higgs model (Higgs and Derrida, 1991; de Aguiar, 2017). The offspring inherits, *locus* by *locus*, the allele of either parent with equal probability, but each *locus* is also subjected to mutations with probability *µ* (*µ* = 0.00025 in our simulations). Offspring are either positioned at the location of the focal parental or it disperses with probability *D* (here we set *D* = 0.01) to one of the 20 nearest sites. This guarantees that close to the location of every individual of the previous generation there will be an individual in the present generation, avoiding the formation of spatial inhomogeneities. The process is repeated until all *M* individuals had a chance to reproduce, completing one generation.

If the number of individuals in the mating range *S* is smaller than a threshold value *I* (*I* = 3 in our simulations) the individual expands the search radius to *S* + 1. If the number of compatible mates is still smaller than *I*, the process is repeated once more to *S* + 2, and if there is still less than *I* potential mates, another neighbor is randomly selected to reproduce in its place. If the neighborhood is empty, which might happen for small values of *S*, reproduction fails and the population decreases by 1. In that case individuals of the next generation will have the opportunity to have a second offspring in order to increase the population back to the specified value *M*. Therefore, for small mating neighborhoods the population size might fluctuate slightly around *M*.

Species are identified as groups of individuals reproductively isolated from all others by the genetic threshold *G* (de Aguiar et al., 2009; Costa et al., 2019). It may happen that two individuals of the same species are above the genetic threshold but have an indirect gene flow through a third individual, which can mate with both. No condition, such as the closest individual in spatial distance or with higher genetic similarity for mating, is imposed on the members of a species. At the beginning, individuals are genetically identical and uniformly distributed in the space.

## Evolutionary events and their detection

Throughout the evolutionary process, mutations and recombinations create genetic diversity that will eventually lead to speciation for certain sets of parameters (de Aguiar et al., 2009; Costa et al., 2019). Because there is a probability *Q* of non-reproduction associated to each individual, the number of species does not remain constant in time. Extinction and hybridization naturally occur during evolution and are, therefore, emergent processes in our model. To detect and characterize these events, we define them in detail below. They are also illustrated in Figure 1.

**Figure 1:**
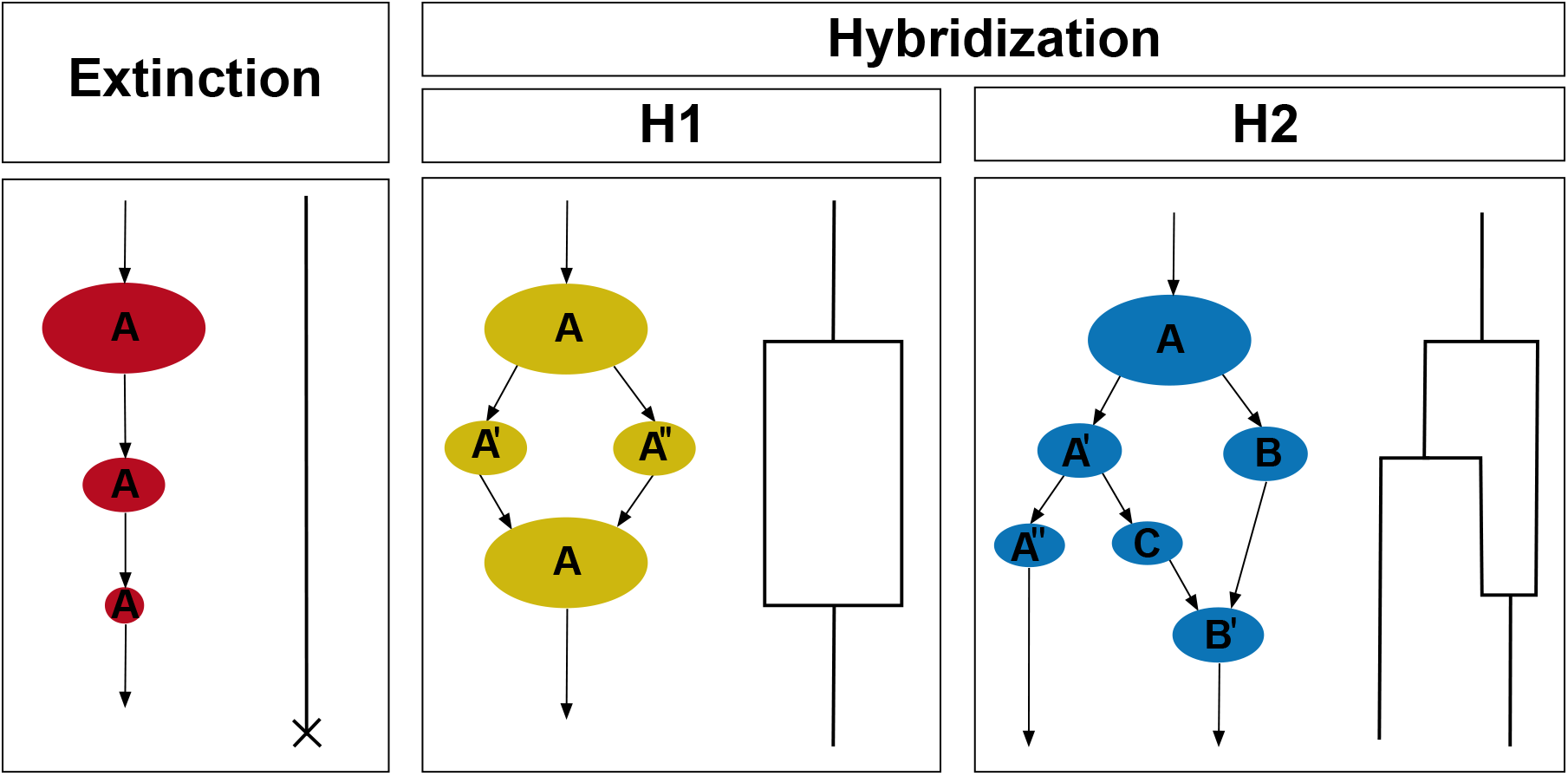
Representation of species population sizes and the respective phylogenetic tree scheme for each event. A true extinction requires that the species size decreases until no descendants are left for the next generation. An H1 event requires that two species (here *A*′ and *A*″) hybridize after one speciation process. An H2 event requires that two species (here *C* and *B*) hybridize after at least two speciation events.

### True extinctions

Since not all individuals reproduce, there are random fluctuations in species abundances that might lead to true extinction. True extinctions will be identified as an event in which individuals of a species leave no offspring to the next generation. This happens when the species has a very small number of individuals (usually of the order of 3 or less) and none reproduces either by chance (probability *Q*^3^ ≈ 0.05 for a species with 3 individuals) or by lack of compatible partners in their mating neighborhoods.

### Hybridization after one speciation process – H1

An H1 event occurs when a species *A* splits into *A*′ and *A*″ and these new species merge back into a single species, as illustrated in the second panel of Fig. 1. Right after speciation, the genetic distance between pairs of individuals belonging to *A*′ and *A*″ is only slightly larger than the threshold *G*, and a few mutations (usually a single one), as well as genetic recombination, might make *d*^(*i,j*)^ ≤ *G* for some *i* ∈ *A*′ and *j* ∈ *A*″. This phenomenon is more probable for small genome sizes. For large genomes, the chance that a mutation will happen in the right locus to decrease the genetic distance between individuals, for example, is small.

### Hybridization after two speciation processes – H2

An H2 event occurs when a species *A* splits into *A*′ and *B* and *A*′, in turn, splits later on into *A*″ and *C* (see third panel of Fig. 1). An H2 event occurs if either *A*″ or *C* merges with *B*. What is remarkable about this phenomenon is that *A*″ and *C* have a more recent common ancestor than they have with *B*, but instead of merging with each other, one of them merges with a more distant species.

### Branch length and population sizes

In the simulations, we kept track of population sizes at the time the branches were interrupted by a true extinction or by a hybridization event. We also kept track of branch lengths, in all cases, to understand the distribution of times of species until extinction and hybridization events. To compute the population size and branch length, we considered the species with the smallest population size in each hybridization event. The largest species was considered the extant one at the moment of the event.

### Probability of hybridization

To estimate the probability of hybridization, we consider two recently diverged species *S*_1_ and *S*_2_ and calculate the probability that offspring, belonging to these species, will be able to mate after mutations and genetic recombination of their parents’ genomes. Mating will restore gene flow and, therefore, merge the species together.

In our model, the minimum genetic distance between parents in *S*_1_ and *S*_2_ is *G*. Let the average genetic distance (normalized to genome size) be (*G*+Δ)*/B* ≡ *g*+*δ*. Defining the similarity between individuals *i* and *j* by *q*^(*i,j*)^ = 1 − 2*d*^(*i,j*)^*/B*, the minimum similarity required for mating is *q*_*min*_ = 1 − 2*g* and the average similarity between individuals in *S*_1_ and *S*_2_ is *q*_*δ*_ = 1 − 2(*G* + Δ)*/B* = *q*_*m*_ − 2*δ*. The distribution of similarities *ρ*(*q*) between offspring from *S*_1_ and *S*_2_ can be approximated by a Gaussian with average *q*_*δ*_ and variance 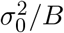, where 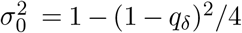 (see SM). The probability that offspring will reinstate the genetic flow between *S*_1_ and *S*_2_ is

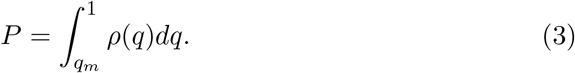

In section 3 of the Supplemental Material, we shown that *P* depends critically on genome size and is given by :

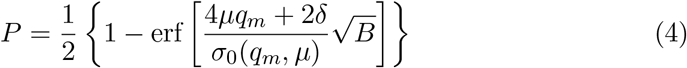

where erf(*x*) is the error function: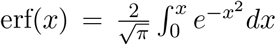. As genome size *B* increases, the error function approaches 1 and hybridization events stop. Because the argument involves the square root of *B*, we expect an exponential decrease in the probability of hybridization with genome size.

## Results

Here we present results of simulations for different genome sizes (*B* = 150, *B* = 1, 500 and *B* = 15, 000). In all cases we ran 50 simulations for 5,000 generations. Figure 2 shows how the simulations look like. The colorful plots represent snapshots of the spatial community at a given generation. The small squares and stars represent the individuals and the different colors represent the different species. The first row highlights a species undergoing a true extinction process, with its individuals depicted by red stars and decreasing in number; at the last temporal plot, only one individual of this species is still alive. Because no reproduction is possible, for lack of sexual partners, this species is extinct in the next generation. The second row represents an H1 event: the individuals of the species represented by black stars split into two species (black stars and yellow stars) and after some generations, the individuals represented by yellow stars hybridize with the individuals represented by black stars, forming only one species again. An H2 event is represented on the third row, in which an initial species (represented by black stars individuals) splits twice – into the species represented by the individuals in blue stars and after that, into the species represented by the individuals in orange stars. After the second split, individuals of the species which splitted first (blue stars) hybridize with the original species (black stars). The full phylogenetic tree containing all the example events (represented by the stars) and also other events of true extinctions (red circles), H1 (yellow circles), and H2 (blue circles), as well as extant species, represented by green circles. In the Supplementary Material (SM), in Fig. S2, we show the same phylogenetic tree of Fig. 2, with only the extant species, as usually phylogenetic trees are represented.

**Figure 2:**
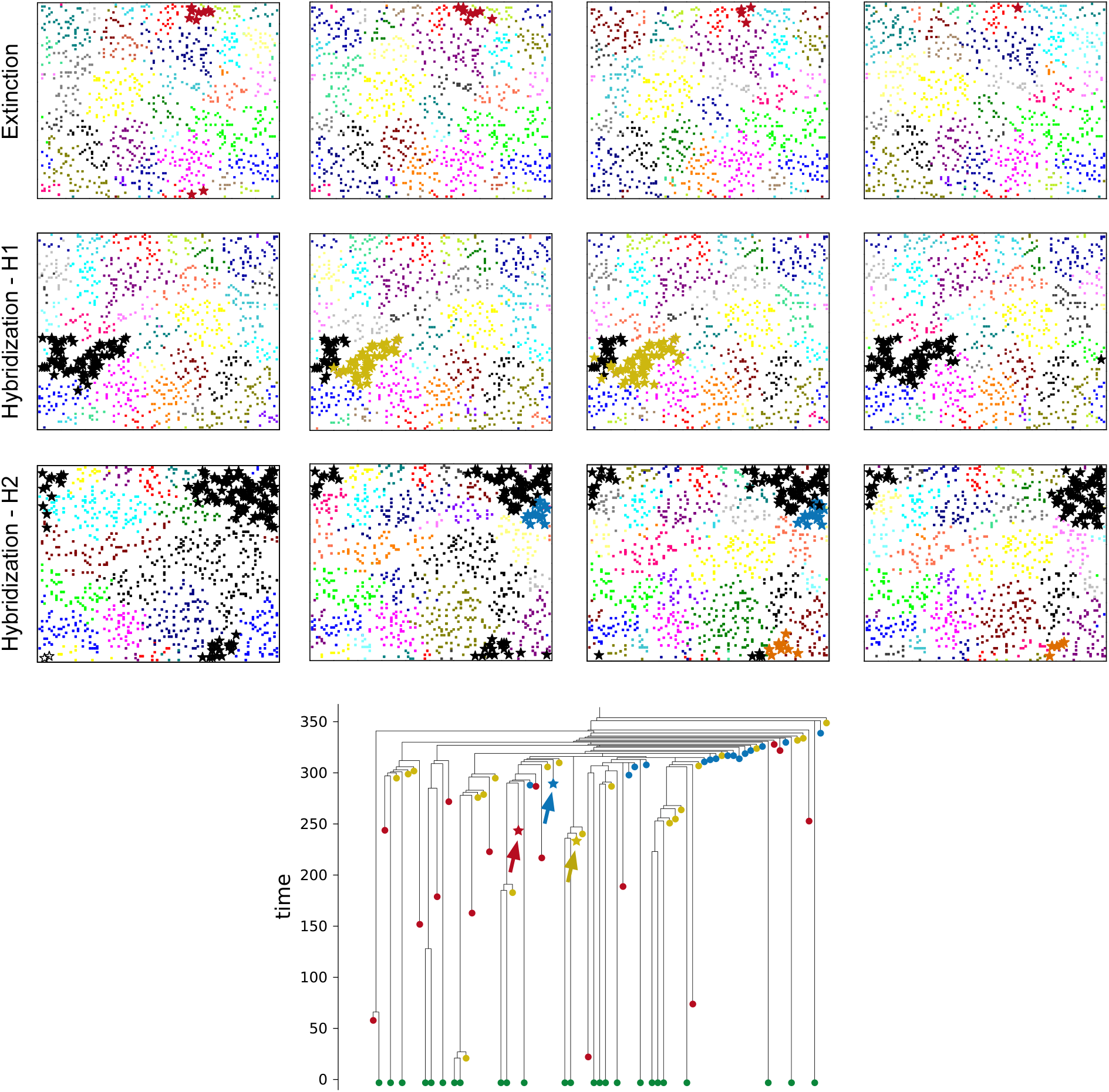
Representation of the three events (true extinction, H1 and H2) in a temporal sequence (above) and in the full phylogenetic tree of events (bellow). The exemplified events are represented in the full phylogenetic tree by stars (true extinction in red, H1 in yellow and H2 in blue) and pointed by arrows. The full phylogenetic tree also has all the other events of true extinctions (red circles), H1 (yellow circles), and H2 (blue circles). The extant species are represented in green circles. See text (Results Section) for more information.

The cumulative number of events along 5,000 generations, for 50 simulations, are represented in Fig. 3. True extinctions are more frequent for larger genomes (*B* = 1, 500 and *B* = 15, 000) and happen under a constant rate for all genome sizes. Hybridization processes are more frequent for smaller genome and have an accelerated rate in the beginning of the evolutionary process. H1 events becomes constant after some time (approximately after 1,300 generations for *B* = 150 and as early as 100 generations for the larger genomes); H2 events, on the other hand, vanish after some generations, reaching a null rate after an initial positive start.

**Figure 3:**
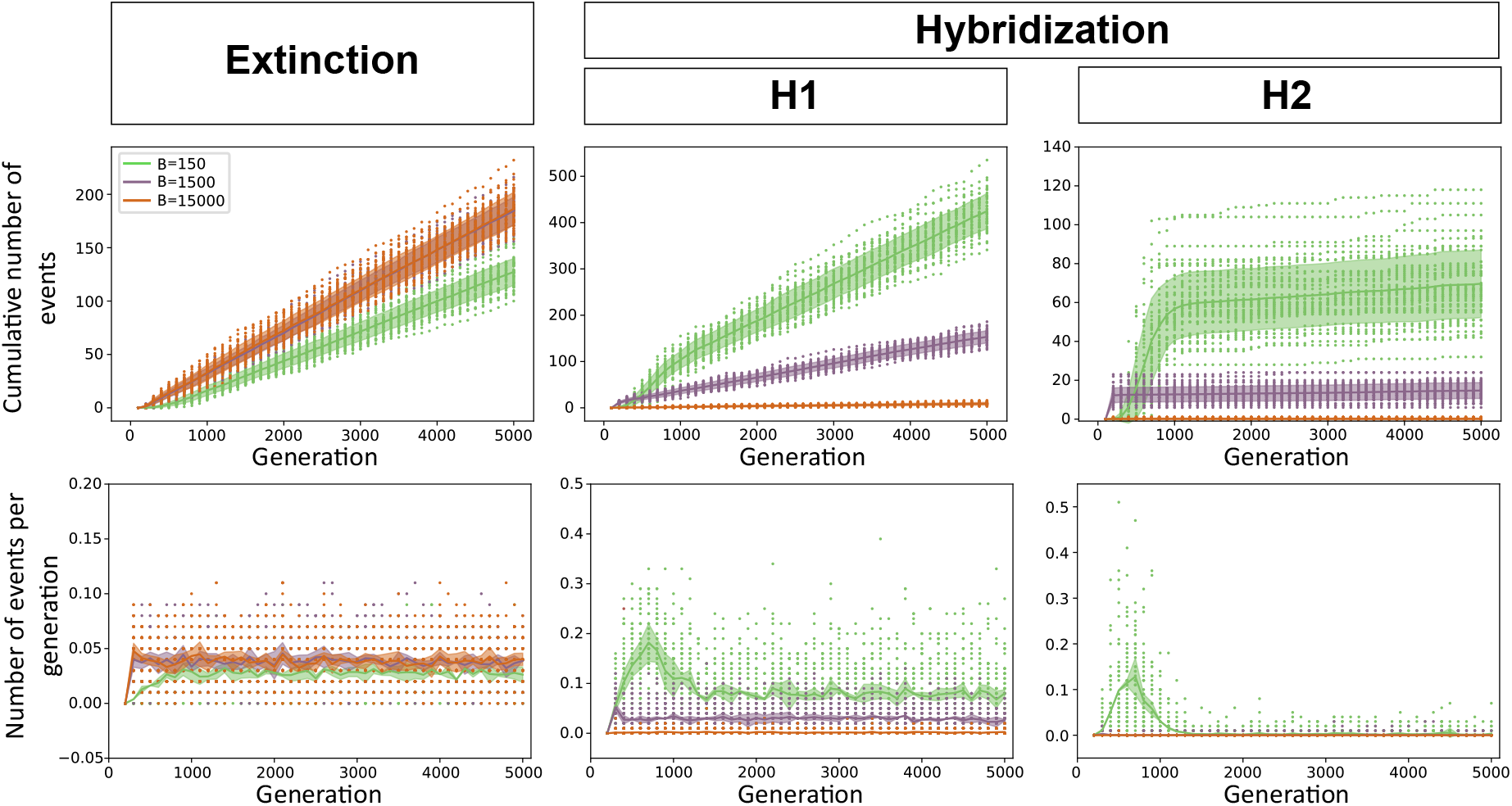
Cumulative number of events along 5,000 generations for 50 simulations (first row) and the respective rates (second row) for three different genome sizes (*B*). Colored points represent simulations, lines are averages over simulations and shaded area are standard deviations.

The population size at the moment the event happened varies according to the event (Fig 4). True extinctions always present low population sizes, measured at the last generation the species existed, which is consistent to how we define it. In average, species reach a population size of 2 individuals before vanishing by true extinction. Hybridization events happen in larger population sizes, with an average of 50 individuals, which is about the average size of extant species after the species richness equilibration for the parameters used in the simulation (see Fig. S5 in the SM). Similar distributions for the population sizes are observed for the other genome sizes for each event (see results for *B* = 1, 500 and *B* = 15, 000 in Fig S3 of the SM).

**Figure 4:**
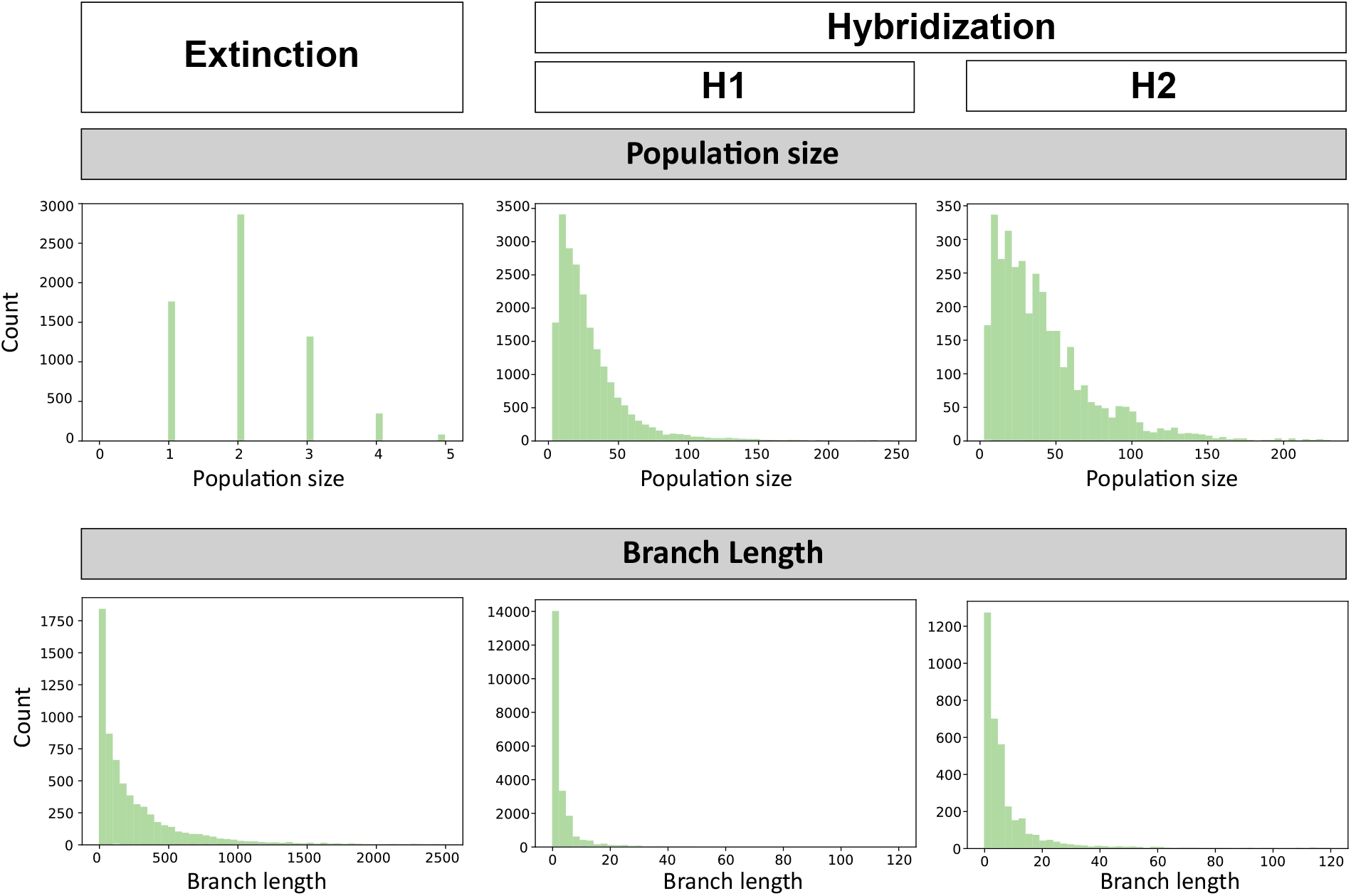
On the first row, population size at the moment of each event for *B* = 150 (see Fig. S3 and text for more details for *B* = 1, 500 and *B* = 15, 000). On the second row, distribution of the branch length from the speciation moment until each event moment for *B* = 150 (see Fig. S4 and text for more details for *B* = 1, 500 and *B* = 15, 000). Results were accumulated along 5,000 generations for 50 simulations.

Species duration, measured by the branch length of the species from the moment it appeared until the moment of the event, is different for each type of event (Fig 4). True extinctions happen both in recently formed species (short branch lengths) and in old species, with a long history (long branch lengths). Hybridization events, on the other hand, always have short duration, *i*.*e*. H1 and H2 events happen few generations after the species branching. H1 events occur typically 10 generations after the splitting, whereas H2 events have a significant probability of occurring later, after 20 generations. These differences between the two mechanisms of hybridization are expected, since H2 events require at least two speciation events, whereas H1 events may happen after only one speciation event. The results for different genomes sizes are presented in Fig S4, in the SM. H1 events are always more frequent than H2 events, but for larger genome sizes, hybridization events become more rare. True extinctions have similar distributions for all genome sizes.

Fig. 5 shows the probability of hybridization – *P* (*B*) – according to the theoretical model (continuous line) for each genome size in relation to the genome size *B* = 150. We present the average (black circles) and error bars of frequencies observed in the simulations. The events of hybridization were counted at the end of generation 5,000. Notice that hybridization events for genomes larger than *B* = 1, 500 are less than 50% of those observed for *B* = 150, and also that fluctuation around the average is greater for smaller genomes.

**Figure 5:**
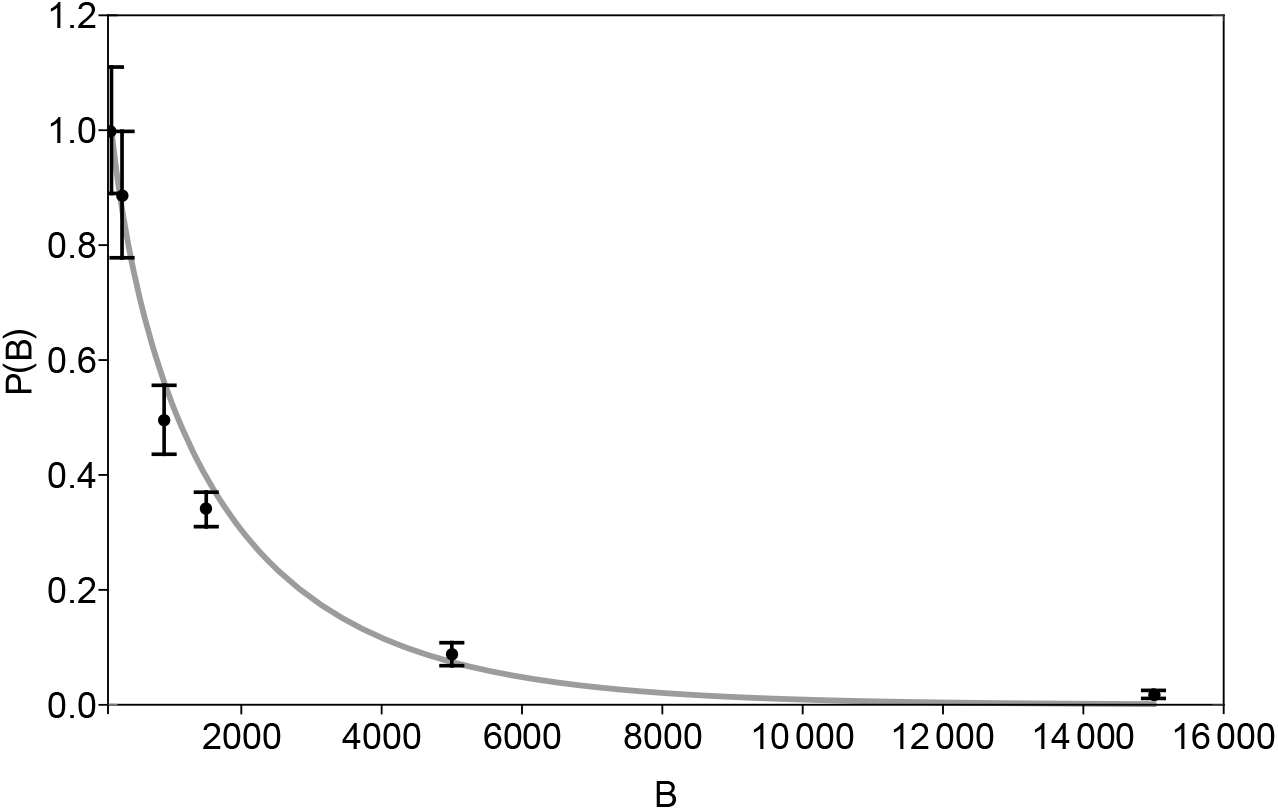
Probability of hybridization *P* (*B*) as a function of genome size *B*. Line shows the theoretical prediction and black circles with error bars show the results of simulations. Points were scaled so that *P* (150) = 1.

Fig. 6 shows the number of successful speciation events (SSE) by the total number of speciation events (SE) accumulated over 100 generations and grouped, by colors, every 500 generations. We define SSE in a time interval as the difference between the number of extant species at the end and beginning of the interval, added to the number of true extinctions that happened in that same period of time. For example, if there are 10 extant species at t=1000 and 12 at t=1100, SSE would be 2 if there were no extinctions. If 1 true extinction is registered, then another successful speciation event must have occurred and SSE=3. The total number of speciation events – SE, is the sum of SSE with the number of hybridization events. We do not consider hybridization event as SSE, since its net effect is zero. Fig. 6 shows that smaller genome sizes have larger number of hybridization events than larger genomes. Moreover, the radiation period for larger genomes, included in the period of 0 to 500 generations (see Fig. S1 in the SM), presents a higher number of events compared to the equilibrium, what is evidenced by the separation of the points in the plot.

**Figure 6:**
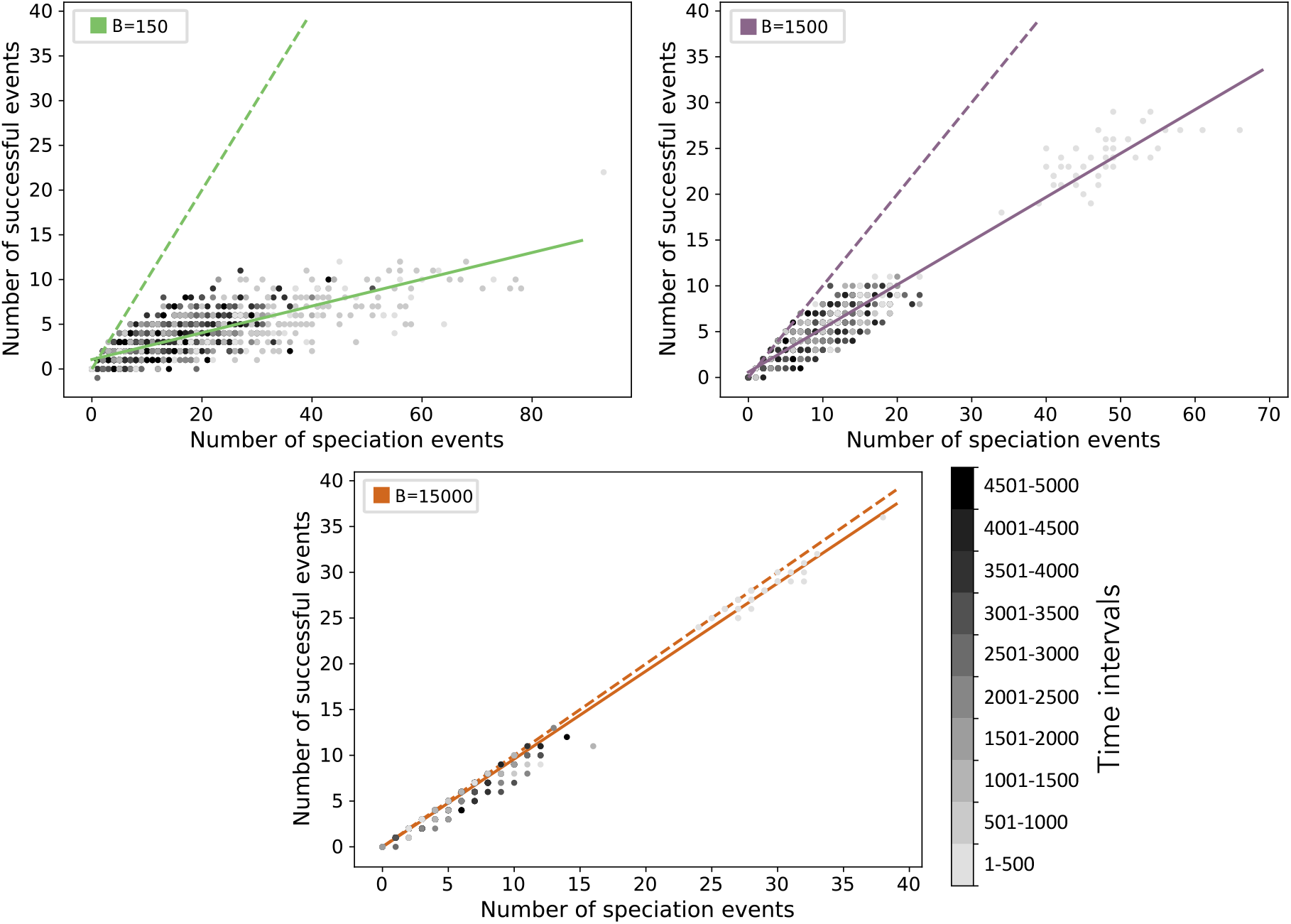
Number of successful events by the number of speciation events. We accumulate events every 100 generations. The plotted points are colored according to the interval in the gray scale. Therefore, there are 5 points at each of these intervals. Genome sizes are *B* = 150 (top left), *B* = 1500 (top right) and *B* = 15000 (bottom). The dashed line is the 1:1 relationship and the continuous line is the regression given by the points of simulations.

## Discussion

In this paper we simulated the process of speciation under a neutral evolutionary scenario using an individual based model. The information stored during the evolutionary process allowed us to reconstruct complete phylogenetic trees, containing information, not only about the living species and their histories, but also about the species that disappeared along the process. We classified the disappearance of species into two basic types of events: true extinctions and hybridization, which were then further subdivided into hybridization after one speciation event, H1, and hybridization after two speciation events, H2. We analyzed how these events differ according to (i) when they occur in the evolutionary history, computing rates of occurrence along the time; (ii) species abundances at the moment of the event and; (iii) the duration of the species up to the moment of the event. All these features were strongly affected by the genome size, which can be interpreted as the amount of information, or traits, that are important for the genetic isolation and therefore to speciation.

Extinctions are hard to infer in the absence of detailed fossil record, but molecular methods to estimate extinction rates can be promising, although in some cases they are still controversial (Rabosky, 2010; He and Hubbell, 2011). Developing methods to solve the problem of underestimation of the number of extinction events is an active field of research (Pimm et al., 2014) and there are numerous evidences that population size is a crucial factor to the extinction of a species (Matthies et al., 2004; O’Grady et al., 2004; Stanley, 1986). In our work, the gradual decrease of the population size until no descendants are left to the next generation is called a true extinction. Since our model is neutral, true extinction events are driven by ecological drift, *i*.*e*., random fluctuations of populations sizes.

We found that genome size affected the rate of true extinctions, with larger genomes leading to slightly higher rates, specially during the radiation. We characterized how long species lasted until they disappeared by true extinction and we found that the distribution of species lifetimes followed the distribution of branch lengths of the extant species. This is expected since there are no forces or processes, other than stochastic fluctuations, that can increase the probability of a species face true extinction. Ecological drift is believed to be of great importance in non-neutral communities (Orrock and Watling, 2010; Gilbert and Levine, 2017). Here we offer a paradigm of how ecological drift on a neutral evolutionary model can be useful as an benchmark for comparison with extinctions in non-neutral models. For instance, the effect of human disturbance on extinction rate of several species has been measured but lacks an expectation for comparison (Pimm et al., 2014). Although extinction estimation is a recurrent theme in neutral models in ecology and evolution (Lande, 1992; Pannell and Charlesworth, 1999; Hubbell, 2001), we believe this is the first time this estimation was made under a macroevolutionary perspective of a microevolutionary model.

Hybridization changes the tree-like structure of phylogenies into more complex, reticulated evolutionary histories. Known factors that promote hybridization include environmental conditions, such as water turbidity (Seehausen et al., 1997; Taylor et al., 2006) and climate change (Seehausen et al., 2008); ecological interactions as parasites-mediated selection for hybrids (Kleindorfer et al., 2014); misprinting (Grant and Grant, 1997, 2014) and the scarcity of sexual mates (Willis et al., 2011). We showed that hybridization (both H1 and H2) also occurs in the absence of selective forces and are more frequent if the genome size is small, decreasing as the genome size increases and reaching nearly zero for *B* = 15000, in accordance with Eq. 4. This is clearly evidenced by the relationship between the number of successful events and the number of speciation events for the different genomes sizes, which is shown in Fig. 6, where the difference between the estimated linear regression and the slope 1:1 is the average number of hybridization events and can also be observed in Fig. 3, in which it is shown the cumulative number and the respective rate of each event, for each genome size.

Hybridization is related to rapid speciation (Lamichhaney et al., 2018), *i*.*e*., to the radiation period, in which the number of extant species is still rising. We found that H2 are rare events and mainly occur during the initial radiation process, while H1 are more common and happens throughout the whole evolutionary process. The probability that mutations and recombination will lead to an H2 event between two species is very small since it requires at least two speciation events, which takes several generations. On the other hand, if the speciation process is recent, H1 is more likely, as the species have more similar genomes. Therefore, H1 events of recently split species are expected to happen during the whole evolutionary history, being responsible for a large proportion of the hybridization events, while H2 events are expected to happen mainly during the radiation, when species still have similar genomes. This is shown in Fig. 6 where it is possible to observe a clear separation between the radiation process (light gray points) and after species richness equilibration (darker points) for genome sizes *B* = 1500 and *B* = 15000. For the smallest genome size (*B* = 150), this separation is not as clear. However it is still possible to identify a more separated group of light gray points, which can be assigned to H2, since they are detected mainly during the radiation, and rarely occur for larger genomes.

The analysis of population sizes at the moment of the event revealed that the distribution of population sizes for hybridization after one or two speciation processes (H1 and H2) displays a large average abundance, of the order of 50 individuals. Therefore hybridization events are not related to the lack of sexual partners or small population sizes in our neutral scenario: it suffices that genetic flow is reestablished between two individuals of different species. Therefore we expect that population abundances for both H1 and H2 events should be similar to the distribution of extant species, which is indeed observed (see Fig S5 on the SM).

Finally we characterized how long species last until they disappear by each type of event. Hybridization is more frequent for recent species (branch length lower than 40 generations), which shows that hybridization events usually happen soon after species differentiation. Although most known cases of such events agree with this temporal result (Kearns et al., 2018), there are some cases, apparently more rare, in which species that differentiated millions of years ago, generate viable offspring in nature or in lab experiments (Kearns et al., 2018; Káldy et al., 2020). In nature, where geographical barriers often exist, hybridization is constrained by both genetic and spatial distances, likely increasing the characteristic time of this event. In our model, although no spatial barriers exist, species are localized in space and only nearby species can hybridize. However, right after speciation, the sister species are spatially close (see Fig. 2) and if individuals respect the genetic criteria for reproduction, there is a high probability that a hybridization will in fact happen. This results in short characteristic times. Events of type H2, however, display a wider distribution of branch lengths than type H1 (branch length lower than 8 generations), which may be explained by the number of speciation processes involved in each type of hybridization. Considering the speciation rate is approximately constant after an initial radiation, two speciation events would take a longer time to happen than only one. In conclusion, in a neutral scenario, extinctions are events related to demographic aspects while hybridization events are related to the moment of the evolutionary history of the species.

Future works can explore the effect of natural selection, how environment heterogeneity and dynamics (such as habitat loss and barriers) can affect the observed patterns of extinction and hybridization events under this model framework, in the same lines of Gascuel et al. (2015) and Kagawa and Takimoto (2018). Moreover, hybridization events not necessarily lead to species diversity reduction. In fact, it is common that hybridization detected in nature shows hybrid individuals coexisting with the parental forms in sympatry (Duenez-Guzman et al., 2009; Kagawa and Takimoto, 2018; Marques et al., 2019). Therefore, we believe it might be possible to characterize ancestors and hybrids coexistence by the genetic distribution, detecting clusters of similar genomes corresponding to the merging species and the hybrids, or by community formation in the network of gene flow between individuals. It would also be interesting to know how long a species last after hybridization.

Historically, hybridization has been more studied in plant species, and possibly the lack of techniques for detecting hybridization in animal species, added to historical contingencies (Mayr, 1963), resulted in a biased literature (Schwenk et al., 2008). We believe the same bias exists in theoretical studies, with few studies estimating and characterizing it in the most basic models – but see the important contributions of McCarthy et al. (1995); Buerkle et al. (2000); DuenezGuzman et al. (2009); Kagawa and Takimoto (2018). We hope more theoretical studies, new molecular techniques, and important field studies can emerge and prosper on the frontiers of species definition.

## Data archiving

There is no data associated with this paper.

## Acknowledgements

Suppressed due to the double-blind reviewing process.

## Notes

### Competing Interest Statement

The authors have declared no competing interest.

