## Supplementary Material for "Reticulate evolution in a neutral model: speciation, extinctions, and hybridizations"

#### 1 Number of extant species × Time

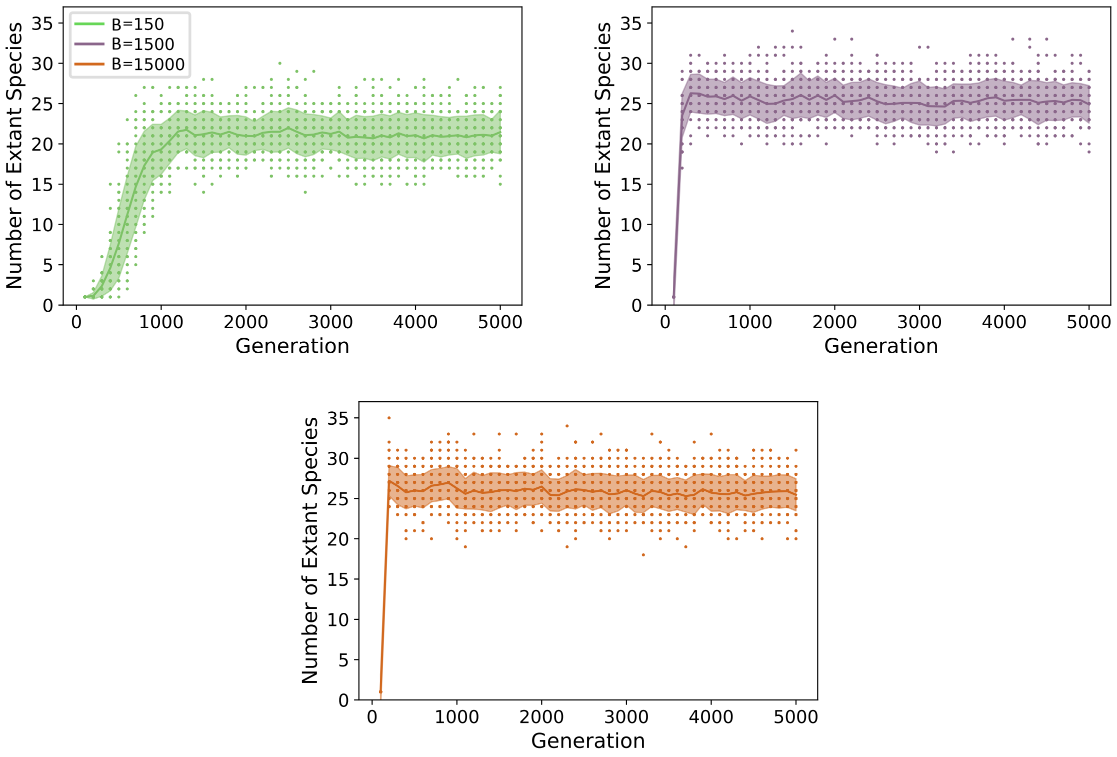


Figure S1: Temporal series of the number of extant species for the different genome sizes. The number of species rise until it reaches an equilibrium, which is between 19 and 24 species for $B = 150$ and between 22 and 27 species for $B = 1,500$ and $B = 15,000$. The equilibration time is reached around 1,100 generations for $B = 150$, around 200 generations for $B = 1,500$ and less than 200 generations for $B = 15,000$.

#### 2 All events tree × Extant species tree

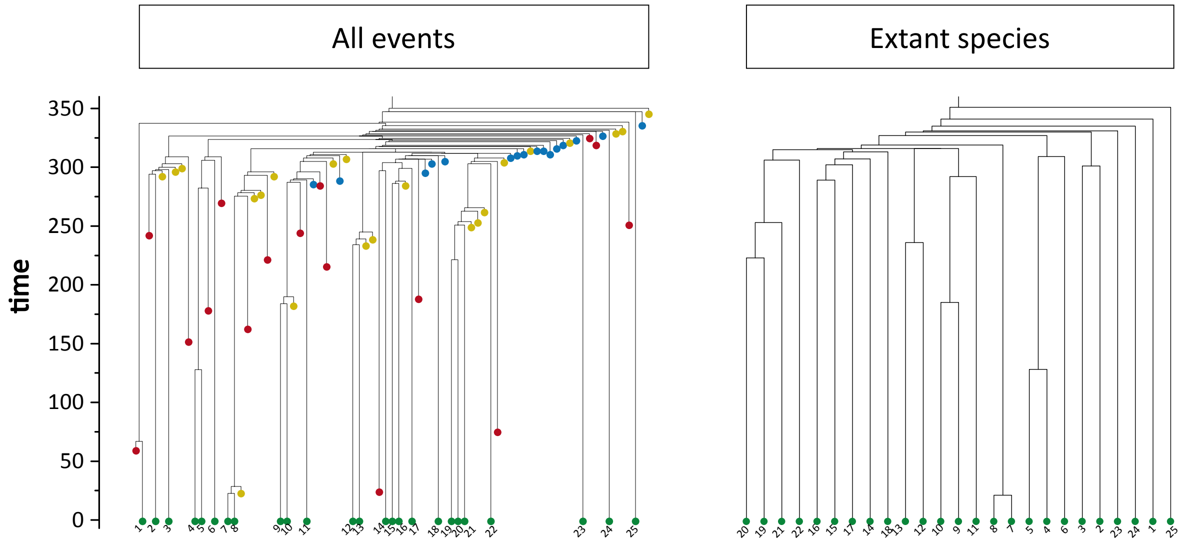


Figure S2: This is the same tree of Figure 2 of the main text. On the left the tree includes all the events (extinctions in red, H1 events in yellow, and H2 events in blue. Extant species are represented in green). On the right we represent the tree by usual phylogenetic tree format, i.e. only with the extant species. The extant species on these trees are not represented exactly in the same order. Species labels are expressed by numbers.

### 3 Complementary information of the distributions of population sizes and branch lengths

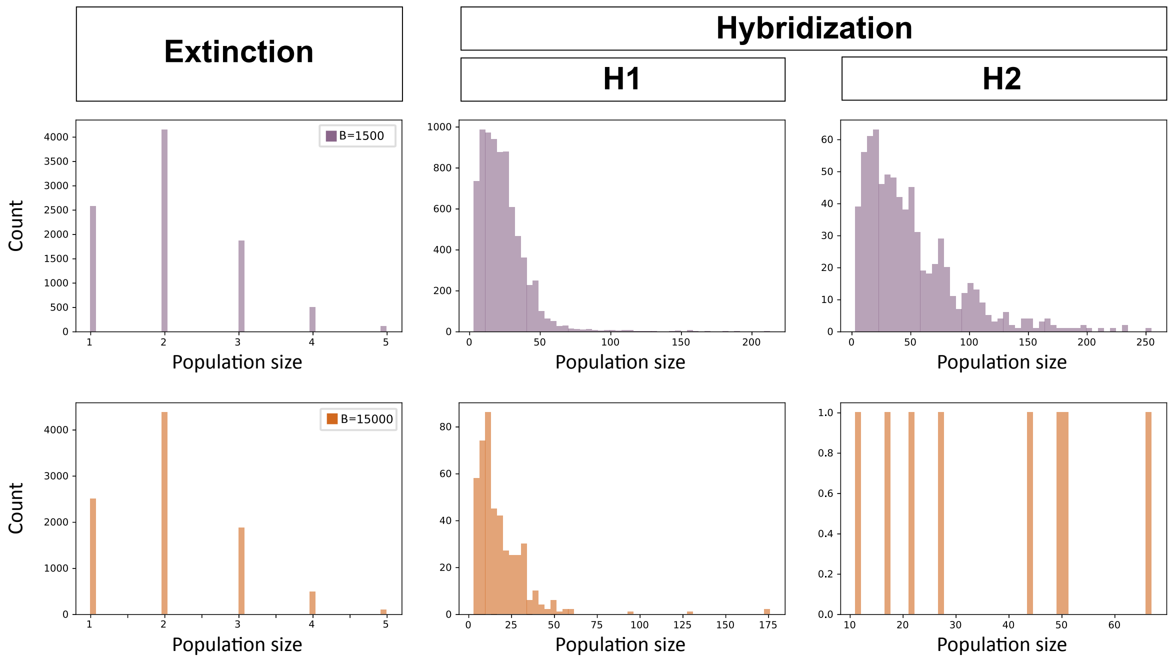


Figure S3: Histograms of the population size at the moment that each event happened, on a linear-linear scale, for $B = 1,500$ (purple, 1st row), and $B = 15,000$ (orange, 2nd row). In the main text, we present the histograms for $B = 150$.

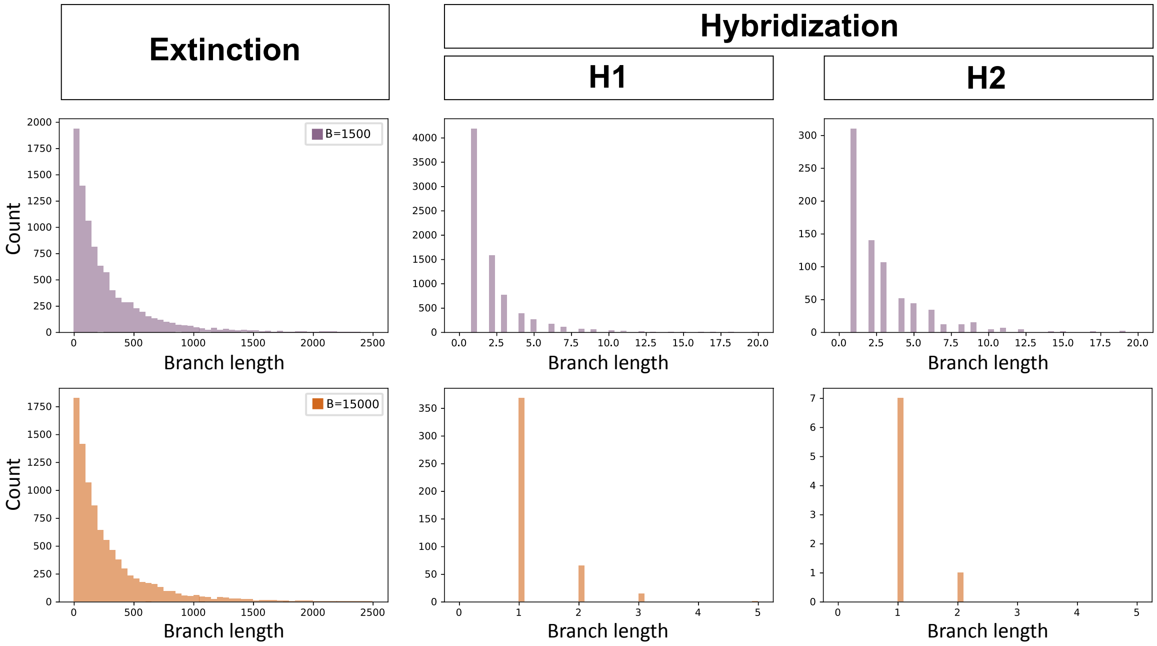


Figure S4: Distributions of the branch lengths from the speciation moment until the moment of each event for $B = 1,500$ (purple, above) and $B = 15,000$ (orange, below). The distributions for $B = 150$ is presented in the main text (Figure 5).

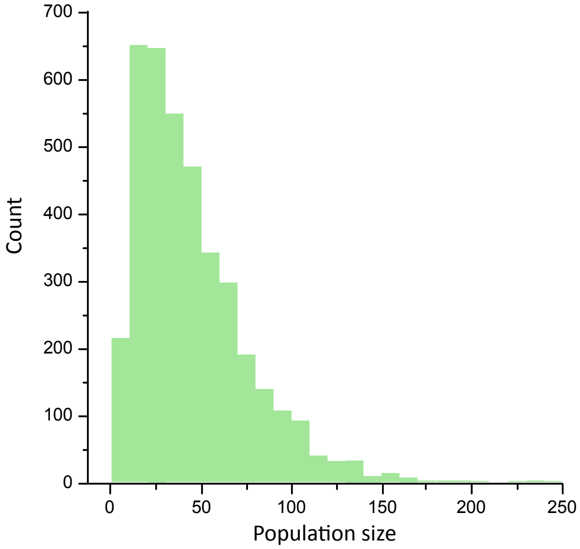


Figure S5: Abundance distribution of the extant species until 1,000 generations for $B = 150$. This histogram follows a log-normal distribution as previously shown by de Aguiar et al. 2009.

### 4 Probability of hybridization events

Suppose two species $S_1$ and $S_2$ have been formed by splitting from a common ancestral species. According to our neutral model, the genetic distance between individuals belonging to $S_1$ and $S_2$ is larger than $G$, the genetic threshold for reproduction. The size of the genome is $B$, and we shall call $g = G/B$ the relative genetic threshold. The average distance between individuals of $S_1$ and $S_2$ is $G + \Delta$, with $\Delta \geq 1$, and the relative distance is $g + \delta$ with $\delta = \Delta/B$.

In terms of similarities we define the minimum similarity for reproduction

$$q_m = 1 - 2G/B = 1 - 2g \tag{1}$$

and also

$$q_\delta = 1 - 2(G + \Delta)/B = 1 - 2(g + \delta) = q_m - 2\delta. \tag{2}$$

Let $P_1(\alpha)$ and $P_2(\alpha)$, both belonging to species $S_1$, be the parents of individual $\alpha$ and $P_1(\beta)$ and $P_2(\beta)$, both belonging to species $S_2$, be the parents of individual $\beta$. We want to know the probability that the genetic distance between $\alpha$ and $\beta$ is smaller or equal to $G$, or equivalently, that their similarity is larger or equal to $q_m$.

We know that the expected value of $q^{\alpha\beta}$ is given by

$$E(q^{\alpha\beta}) = \frac{e^{-4\mu}}{4}\left(q^{P_1(\alpha)P_1(\beta)} + q^{P_2(\alpha)P_1(\beta)} + q^{P_1(\alpha)P_2(\beta)} + q^{P_2(\alpha)P_2(\beta)}\right). \tag{3}$$

Assuming that $q^{P_i(\alpha)P_j(\beta)} = q_\delta$, this simplifies to

$$E(q^{\alpha\beta}) = q_\delta e^{-4\mu} \equiv q_{\delta,\mu}. \tag{4}$$

The variance of this distribution, due to the finite size of the genome, is

$$\sigma^2 = \frac{1}{B} - \frac{e^{-8\mu}}{4B}(1 + q_\delta)^2 \equiv \frac{\sigma_0^2(q_\delta, \mu)}{B}. \tag{5}$$

We will approximate the distribution of similarities by a Gaussian

$$\rho(q) = \frac{1}{\sqrt{2\pi}\sigma} \exp\left[-(q - q_{\delta,\mu})^2/2\sigma^2)\right] \tag{6}$$

and the probability that $\alpha$ and $\beta$ will reinstate the genetic flow between $S_1$ and $S_2$ is

$$P = \int_{q_m}^{1} \rho(q) dq. \tag{7}$$

#### 4.1 Calculation of the probability

Making the change of variables

$$x = (q - q_{\delta,\mu})/\sigma \tag{8}$$

the lower integration limit goes into

$$q_m \rightarrow x_m = [q_m(1 - e^{-4\mu}) + 2\delta e^{-4\mu}]/\sigma \approx [4\mu q_m + 2\delta]\sqrt{B}/\sigma_0. \tag{9}$$

The upper limit may be put to infinity if $q_m$ is not too close to 1 and $\sigma << 1 - q_m$, which is usually the case if $B$ is large. In this case we get

$$P = \frac{1}{\sqrt{2\pi}} \int_{x_m}^{\infty} e^{-x^2/2} dx = \frac{1}{2}[1 - \mathrm{erf}(x_m)] \tag{10}$$

where

$$\mathrm{erf}(x) = \frac{2}{\sqrt{\pi}} \int_0^x e^{-x^2} dx \tag{11}$$

is the error function.

#### 4.2 Interpretation and behavior

The probability of hybridization is

$$P = \frac{1}{2}\left\{1 - \mathrm{erf}\left[\frac{4\mu q_m + 2\delta}{\sigma_0(q_m, \mu)}\sqrt{B}\right]\right\}. \tag{12}$$

The calculation involves the unknown parameter $\delta$ that gives the relative distance between two species right after the split. For fixed $\delta$, $\mu$ and $q_m$ we see that the argument of the error function increases monotonically with $B$ and the probability of hybridization decreases monotonically with $B$, as observed in the simulations.

The probability has two contributions: the term containing $\delta$ is responsible for hybridization by pure recombination, as it survives even if $\mu = 0$, whereas the term proportional to $\mu$ gives the probability of hybridization due to mutations. In Fig. 6 of the main text, we use $\delta = 0.0025$, $q_m = 0.9$ and $\mu = 0.00025$. We fitted Eq. 12 using $B = 150$, as a reference point, dividing the probabilities for all the other values of $B$ by the probability given by $B = 150$.
